# Reconstructing coniferous tree crown shape from incomplete point clouds using deep learning

**DOI:** 10.64898/2026.01.18.700158

**Authors:** Aline Bornand, Meinrad Abegg, Felix Morsdorf, Stefano Puliti, Rasmus Astrup, Nataliia Rehush

**Affiliations:** Forest Resources and Management, Swiss Federal Institute for Forest, Snow and Landscape Research WSL, Zuercherstrasse 111, Birmensdorf, 8903, Switzerland; Department of Geography, University of Zurich, Winterthurerstrasse 190, Zürich, 8057, Switzerland; Division of Forest and Forest Resources, Norwegian Institute for Bioeconomy Research (NIBIO), Høgskoleveien 8, Ås, 1433, Norway

**Keywords:** point cloud completion, deep learning (DL), LiDAR, terrestrial laser scanning (TLS), mobile laser scanning (MLS), forest, trees

## Abstract

Individual tree structure plays a key role in forest monitoring, biomass estimation, and ecological assessment. However, ground-based remote sensing methods such as terrestrial and mobile laser scanning frequently produce incomplete point clouds due to occlusion, particularly in the upper canopy. This limits the accuracy of derived structural metrics such as tree height or crown volume. In this study, we present a novel deep learning-based method to reconstruct the outer crown shape of coniferous trees from incomplete point clouds. Instead of completing the full tree structure, we focus on predicting the alpha-shape of the crown, enabling a more efficient and generalizable approach for structural reconstruction. We train a geometry-aware transformer model (AdaPoinTr) on synthetically generated partial tree crowns and evaluate its performance across three independent datasets encompassing different forest types and acquisition conditions. The model consistently improved crown shape similarity metrics and reduced height estimation errors compared to using partial data alone (reduced bias from -11% to -3.5%). Our results demonstrate that this shape-based strategy enables the extraction of key tree-level parameters from incomplete data, offering a practical solution for gaining improved 3D forest structural information from cost-sensitive or logistically constrained forest monitoring acquisitions.

## 1. Introduction

Structural information of individual trees is essential for forest monitoring and ecosystem research. Metrics such as tree height, crown dimensions, and crown volume are crucial for estimating biomass, assessing competition, and quantifying habitat complexity and biodiversity (Jucker et al., 2025). These parameters support both forest management practices and ecological studies aiming to understand forest dynamics and ecosystem functions.

Three-dimensional forest structure is increasingly characterized using light detection and ranging (LiDAR) technology. Airborne laser scanning (ALS) offers extensive spatial coverage, whereas close-range methods—including terrestrial laser scanning (TLS), mobile laser scanning (MLS), and unoccupied aerial vehicle laser scanning (UAVLS)—enable high-resolution measurements of individual trees due to their dense point clouds. These close-range techniques support detailed, non-destructive assessments of crown morphology (Kunz et al., 2019), vegetation density (Grau et al., 2017), and above-ground volume and biomass (Demol et al., 2022; Brede et al., 2022b).

However, capturing complete point clouds in forest environments is often hindered by occlusion from branches, foliage, and neighbouring trees (Kükenbrink et al., 2017). Scanner positioning, limited range, and environmental complexity frequently result in missing data. This limits the accuracy of tree architecture reconstruction and volume estimation (Morhart et al., 2024), and even basic measurements such as tree height (Wang et al., 2019). While strategies such as multi-scan TLS setups (Abegg et al., 2017), and data fusion with ALS or UAVLS (Giannetti et al., 2018; Terryn et al., 2022) can reduce occlusion effects, these approaches may be impractical in rugged terrain, dense canopies, or due to cost constraints. When scanning from the ground using TLS or MLS, incomplete or missing crown sections, especially in the upper canopy, are a recurring challenge. Consequently, TLS often underestimates tree height due to the difficulty of capturing treetops obscured by foliage or neighbouring trees (Cabo et al., 2018; Vaglio Laurin et al., 2019; Wang et al., 2019; Liang et al., 2018). Mathes et al. (2023) observed this effect at both the individual and stand level, noting that height underestimation was more pronounced in spruce than in beech, and that substantial information was lost in the canopy when relying solely on ground-based scanning. This limitation persists even when advanced scan setups are used, as noted by Hämmerle et al. (2017), who found that occlusion-induced underestimation remains a challenge despite simulated improvements in scanner coverage.

Similar challenges arise with other ground-based approaches. Terrestrial photogrammetry based on structure-from-motion (SfM) offers a cost-effective alternative for generating 3D forest point clouds, but has a more limited range and capacity for canopy penetration. Ground-based SfM point clouds typically only capture the lower sections of trees, with canopy reconstruction largely absent or inaccurate (Piermattei et al., 2019).

Moreover, even when forest-wide point clouds are complete, subsequent processing steps can still lead to data loss. In particular, single tree segmentation can result in structural incompleteness. Although automated instance segmentation methods have improved considerably (Wielgosz et al., 2024), segmentation errors remain common in dense and complex canopies. In some cases, segmentation boundaries may be too conservative, excluding lateral crown parts and thus causing incomplete single trees (Cherlet et al., 2024).

A promising avenue to address incomplete 3D data during post-processing is the application of point cloud completion techniques based on deep learning (DL). These methods, initially developed in 3D computer vision, have advanced rapidly in recent years since the introduction of PointNet++ (Qi et al., 2017), which enabled the direct processing of raw 3D point cloud coordinates. Building on this foundation, a variety of deep learning architectures have emerged, including convolution-based, folding-based, graph-based, and more recently, transformer-based methods (Fei et al., 2022). The Transformer architecture, originally developed for natural language processing, typically includes an encoder–decoder structure and is now at the core of many state-of-the-art methods for 3D completion tasks (Yu et al., 2023). Although these techniques are increasingly applied across diverse domains, most current deep learning research continues to focus on geometrically regular objects found in indoor or urban environments (Chang et al., 2015), with relatively limited work addressing the irregular, highly variable structures of natural objects such as trees.

We previously demonstrated the potential of DL-based point cloud completion for filling gaps in detailed tree structure point clouds, provided that appropriate training data are available (Bornand et al., 2024). However, most existing completion networks are constrained by computational costs, limiting the number of points per sample. The few existing studies specifically targeting full individual trees (Xu et al., 2025; Zhang et al., 2025; Ge et al., 2024) used relatively sparse point clouds. For example, TreegrowNet (Zhang et al., 2025) was trained to complete individual trees to just 8192 points. Current networks generally perform well for samples under 10,000 points, beyond which GPU memory constraints become a limiting factor.

At this level of detail, the inner crown structures of trees are often noisy or missing, making detailed branch reconstruction unfeasible, particularly for evergreen conifers. However, many forestry and ecology applications rely primarily on external structural metrics such as DBH, tree height, and crown dimensions. In contrast to stem diameter and tree height, detailed crown dimensions (such as crown projection area, crown width, crown length, and crown volume) cannot be accurately obtained through traditional field measurements (Lines et al., 2022). However, these attributes can be derived from 3D point clouds and have proved to be essential for estimating individual tree biomass (Vonderach et al., 2018; Menéndez-Miguélez et al., 2023), analysing structural complexity (Das et al., 2025), or assessing competition among trees (Metz et al., 2013).

For obtaining crown dimensions, we do not necessarily need dense TLS data that contains detailed branch structure of the tree. For these purposes, the outer boundary or “hull” of the point cloud is sufficient. This boundary can be computed using alpha-shapes (Edelsbrunner and Mücke, 1994). Alpha-shapes have the advantage that they allow for tighter and more realistic representations of tree crown geometries than simply using convex hulls. Moreover, Bornand et al. (2023) demonstrated a correlation between alpha-shape volume and the volume of small branches. Wang et al. (2024) evaluated various crown volume extraction methods and found that alpha-shape volumes, particularly with alpha values between 0.3 and 0.6, showed the highest correlation with tree biomass.

Therefore, if available point cloud data are relatively sparse and incompletely represent the canopy, as is often the case with ground-based scans, reconstructing the full tree point cloud may not be necessary. Focusing on the outer hull can already yield valuable structural information, particularly for conifers, while also keeping the DL model comparatively more lightweight and less prone to overfitting.

Here, we propose a novel approach that applies DL-based point cloud completion not to the full tree point cloud, but specifically to its outer hull. We evaluate whether deep learning can reconstruct the crown shape of individual conifer trees from partial input data acquired with ground-based laser scanning. We validated our approach on three independent datasets spanning a range of forest types and acquisition conditions. We assess the similarity of the predicted hull to the full tree’s crown shape, compare predicted tree height to manually measured height, and analyse how varying levels of missing upper canopy data affect reconstruction performance.

## 2. Materials and Methods

### 2.1. Data

#### 2.1.1. Training dataset

For training the completion model, we used data from the FOR-species20K dataset Puliti et al. (2025), which is a collection of segmented individual tree point clouds from TLS, MLS, and UAVLS sources collected across the globe. We focus on the four conifer species most abundant in the dataset (*Picea abies, Pinus sylvestris, Pseudotsuga menziesii, Abies alba*), trees of which were scanned in temperate and boreal forests across Europe. After visually checking the data quality and completeness, we selected 1028 single trees.

#### 2.1.2. Independent test datasets

We used three independent datasets to evaluate different aspects of the performance of the completion model:

**hdALS-MLS (NIBIO)**: This dataset was created by fusing MLS data from a ZEB Horizon scanner with dense ALS data acquired using a Riegl VUX-24 sensor mounted on a helicopter. After an automatic tree segmentation with ForAINet (Xiang et al., 2024), we manually selected 80 complete single trees, which we additionally cleaned in CloudCompare v2.13 (Girardeau-Montaut, 2019). The exact species of the selected trees have not been identified in the field, but we assume them to be *Picea abies* and *Pinus sylvestris*.

**SWBM**: This dataset includes 59 single trees representing *Picea abies, Pinus sylvestris, Pseudotsuga menziesii, Abies alba* and *Larix decidua* (Bornand et al., 2023). The trees were scanned using a Leica BLK360 TLS device. Due to dense canopies and the distance to the scanner, the majority of these point clouds lack the upper parts of the tree crowns. However, the height of these trees was manually measured in the field using the hypsometer Vertex IV (Haglöf, Sweden) and is available as reference data.

**pytreedb**: From the individual tree database pytreedb (https://pytreedb.geog.uni-heidelberg.de), we selected 100 single tree point clouds from a dataset by Weiser et al. (2022). The selected trees consist of high-quality TLS data as well as ALS data of *Picea abies, Pinus sylvestris, Pseudotsuga menziesii* and *Abies alba*. We fused the TLS and ALS point clouds to ensure complete representation of the tree tops.

**Table 1:** Overview of the datasets used in this study.

| Dataset | Purpose | Number of trees | Species | Sensor type | Type of reference available | Partial point clouds |
| --- | --- | --- | --- | --- | --- | --- |
| FOR-species20K (Puliti et al., 2025) | train | 1028 | *Picea abies*,<br>*Pinus sylvestris*,<br>*Pseudotsuga menziesii*,<br>*Abies alba* | TLS, MLS, UAVLS | complete trees | simulated |
| pytreedb (Weiser et al., 2022) | test | 100 | *Picea abies*,<br>*Pinus sylvestris*,<br>*Pseudotsuga menziesii*,<br>*Abies alba* | TLS + ALS | complete trees | simulated |
| hdALS-MLS (NIBIO) | test | 81 | *Picea abies*,<br>*Pinus sylvestris* | MLS + dense ALS | complete trees | real-world occlusion |
| SWBM (Bornand et al., 2023) | test | 59 | *Picea abies*,<br>*Pinus sylvestris*,<br>*Pseudotsuga menziesii*,<br>*Abies alba*,<br>*Larix decidua* | TLS | field-measured tree height | real-world occlusion |

### 2.2. Data preparation workflow

We prepared the training data as illustrated in Figure 2. The input consists of point clouds representing full trees, which are first downsampled to contain between 75000 and 110000 points. To simulate partial tree point clouds, the upper part of the tree is removed using the following method: a top layer is defined as the points located above a randomly chosen height threshold, ranging from 75% to 98% of the total tree height. Within this top layer, a random point is selected as the centre of a sphere. The radius of this sphere is randomly assigned a value between 2% and 50% of the tree height. All points inside the sphere are then removed, resulting in the final partial tree point cloud. We repeated this procedure ten times for each tree in the training dataset.

**Figure 1:**
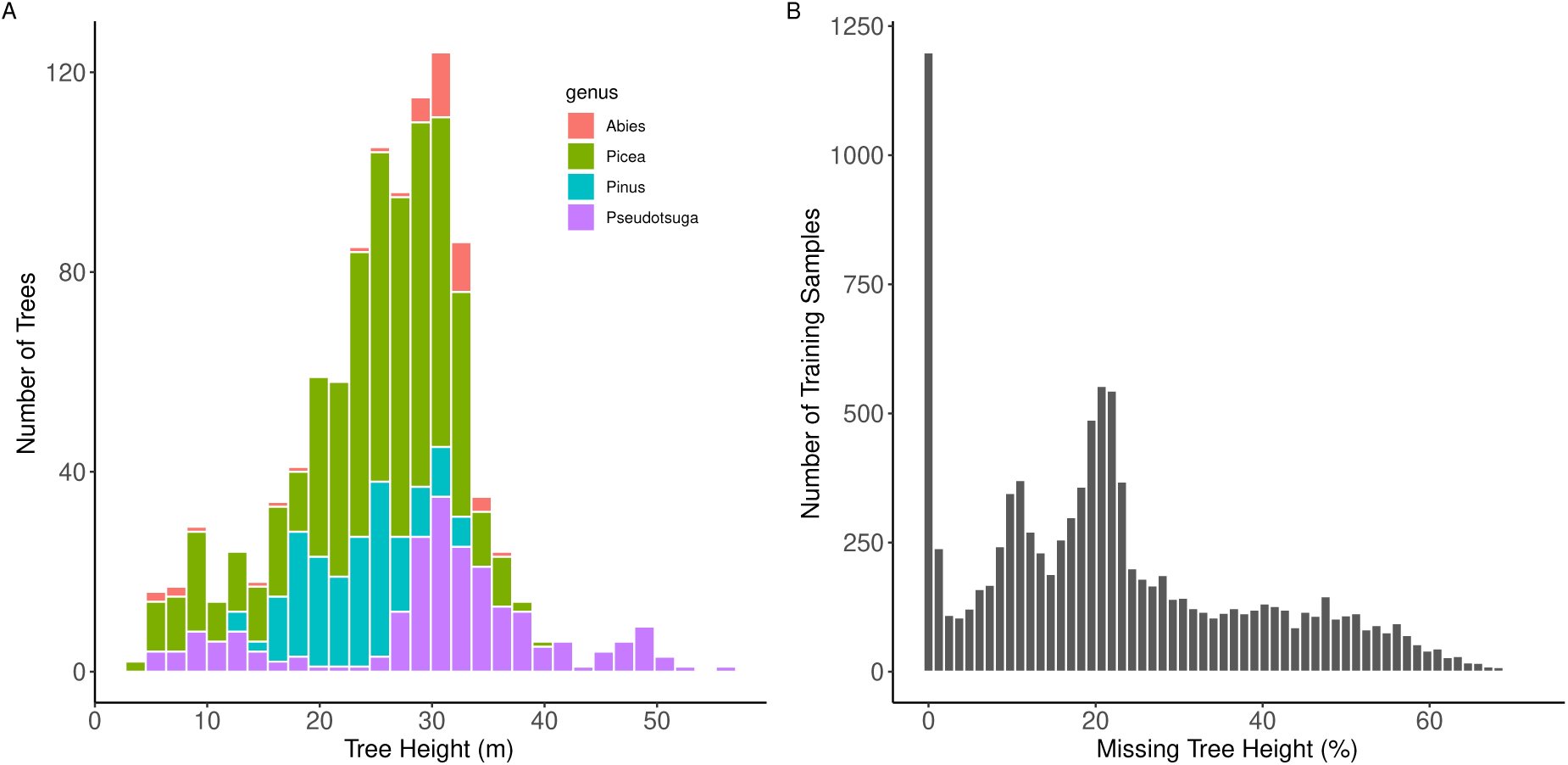
A) Distribution of tree height and genus in the training dataset representing a selection of trees from the FOR-species20K dataset (Puliti et al., 2025). B) Distribution of percentage of missing tree height in the partial samples of the training dataset.

**Figure 2:**
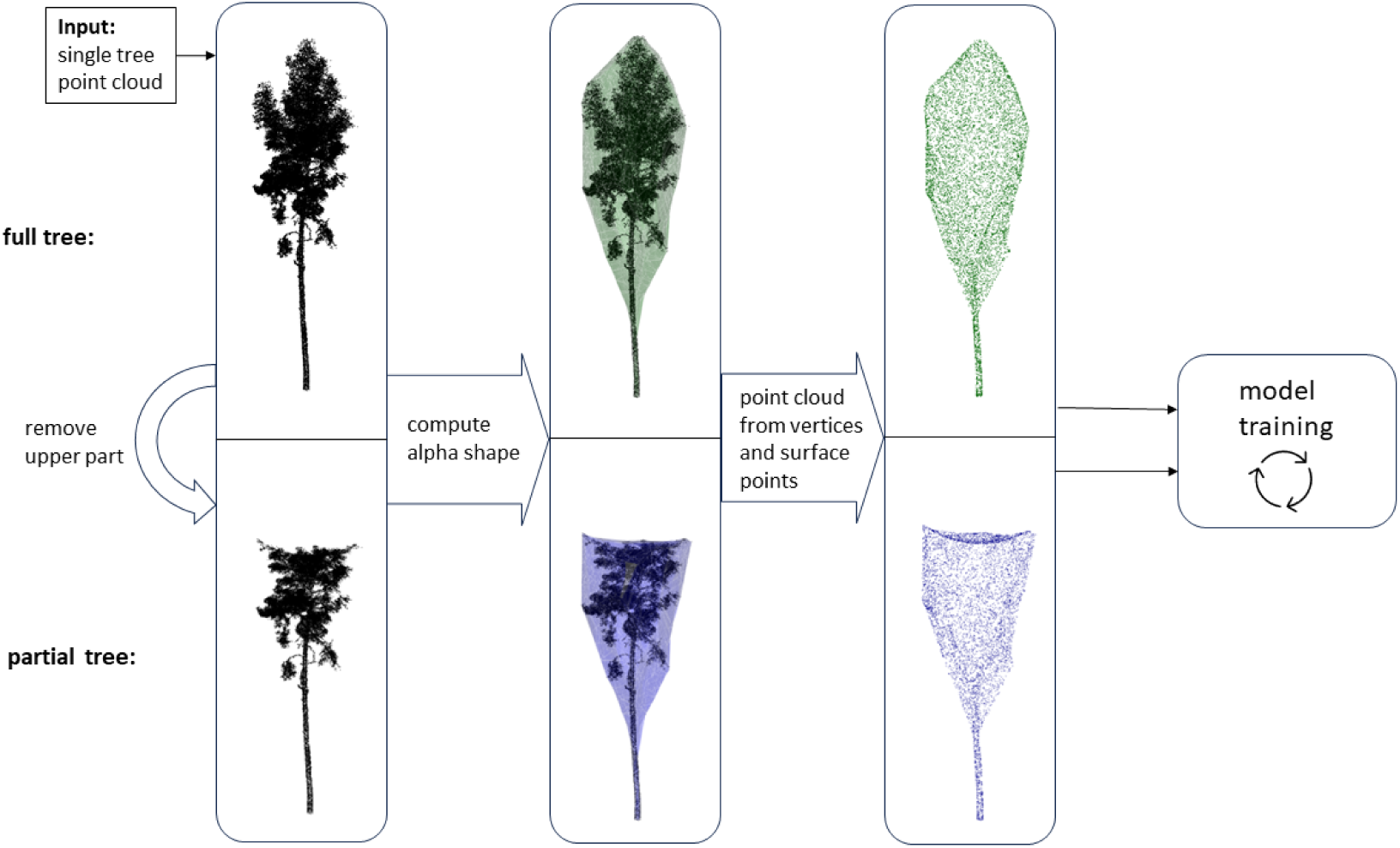
Data preparation workflow: An upper part is removed from the full single tree point cloud, alpha-shapes are computed from the full and partial trees, and alpha-shape vertices are transformed back into point clouds, which then serve as input for model training and testing.

From both the full tree point cloud and the partial tree point cloud, the outer hull is then computed by applying the alpha-shape algorithm using the R-package ’alphashape3d’ (Pateiro-López and Rodríguez-Casal, 2010). The alpha-shape algorithm is a widely used computational geometry method for extracting the boundary of discrete point sets (Edelsbrunner and Mücke, 1994). It operates by conceptually rolling a circle of radius ’alpha’ around the external edges of the point set; when alpha is sufficiently large, the circle cannot enter the interior, thereby tracing the outer boundary of the set. As alpha approaches infinity, the resulting boundary converges to the convex hull of the point set. In the context of tree crown reconstruction, the alpha-shape algorithm provides a more refined representation of the crown boundary compared to the convex hull approach. The choice of alpha-value directly influences the level of detail. Smaller values result in a tighter fit that better captures crown structure but may introduce gaps and unconnected parts, whereas larger values produce a smoother and more convex approximation. For generating training data, we used random alpha-values between 0.25 and 0.34. This range maintains a reasonable balance between shape detail and computational efficiency, resulting in alpha-shapes with fewer than 8,000 vertices, which fits within the 8192-point limit for training samples of the model we used. Smaller alpha values would raise the number of vertices and introduce excessive variation across samples, making the model harder to train and requiring significantly more data and time.

The vertices of the alpha-shape are then saved as a new hull point cloud. If the number of vertices is less than 8192, additional points are randomly sampled on the surface of the alpha-shape mesh to ensure sufficient density. To maintain a continuous surface in partial tree samples, the alpha-shape is computed separately for full and partial tree point clouds. This approach prevents gaps in the partial samples, ensuring that their surface remains consistent, as it would if the hull were computed directly from an already incomplete tree. For the partial samples, the hull point clouds are downsampled to a randomly chosen percentage between 25% and 75% of the total points in the full hull point cloud. This ensures that the model is exposed to a range of point densities during training.

The resulting pairs of point clouds (full and partial hulls) were then normalised (scale and offset) and served as training and test data for the AdaPoinTr model. In total, we generated 10000 data pairs from 1000 individual trees from the FOR-species dataset, meaning that each tree appeared 10 times in the training data with different crown portions removed.

### 2.3. Training of completion model

Transformer models represent the state-of-the-art for generative tasks, including the prediction of 3D points. In the context of point cloud completion, approaches that operate directly on unstructured XYZ data are the most practical and versatile. To complete tree structures, we previously applied and fine-tuned the geometry-aware transformer PoinTr (Yu et al., 2021) on both real and simulated tree data (Bornand et al., 2024). A successor model, AdaPoinTr (Yu et al., 2023), has since been used by Xu et al. (2025); Zhang et al. (2025) as the baseline architecture in methods specifically designed to complete point clouds of full individual trees.

In this study, we also applied the AdaPoinTr model. The AdaPoinTr model includes a geometry-aware Transformer with an encoder–decoder architecture that abstracts the original point cloud into a collection of proxy point clouds, defines global features and then infers the missing proxy points. Compared to PoinTr, AdaPoinTr also integrates an adaptive development query mechanism and a de-noising task, which improves the efficiency and robustness of the model and reduces noise and clumping effects in the predicted point clouds. AdaPoinTr uses Chamfer Distance (CD) as its loss function. CD is the most widely used metric for completion tasks involving unordered point clouds, serving both as an evaluation measure and as a loss function for training learning-based algorithms (Tesema et al., 2023). AdaPoinTr is implemented in Python using the PyTorch library (**?**). Like in many other 3D completion approaches, the AdaPoinTr network is trained and tested on the ShapeNet dataset (Chang et al., 2015), a large-scale open repository of 3D CAD models. The ShapeNetCore collection includes over 51,000 shapes across 55 common object categories. However, these shapes mainly represent artificial objects. Pre-trained model weights based on this dataset are provided by Yu et al. (2023).

Starting from these general weights, we fine-tuned AdaPoinTr on our own training data (see Sections 2.1.1 and 2.2). This fine-tuning approach reduces both the training time and the amount of required data compared to training a deep learning model from scratch.

We trained ten separate models using the same hyperparameters but with different random splits of 80% training and 20% validation data. This strategy allows us to avoid stratifying by species, size, or other factors, while still gaining insight into the variability of predictions across different training/validation splits.

We carried out all experiments on a Linux machine running Ubuntu 22.04.3 LTS. When training on one NVIDIA GeForce RTX 3090 GPU with 24 GB of memory, the models converged (CD reached a plateau) after around 120 epochs and a training time between 40 and 120 hours.

### 2.4. Performance evaluation

To assess the effectiveness of the completion model predictions, we evaluated three key aspects of its performance. First, we examined how well the model can reconstruct the overall crown shape by analysing the similarity between the predicted and reference hull point clouds. Second, we assessed the model’s ability to accurately predict the actual tree height, as measured by traditional inventory methods. Third, we investigated how the proportion of missing tree height influences the quality of the predictions. These evaluations were conducted using the three independent test datasets described in Section 2.1.2.

#### 2.4.1. Crown shape prediction

The MLS data from the hdALS-MLS dataset includes many examples of trees where the highest point of the tree is captured in the point cloud, but one side of the crown is occluded in the MLS data. This type of data allows us to test whether the model can accurately reconstruct the original crown shape in situations where side branches are either occluded from the scanner’s view or missing due to segmentation errors.

We used the MLS point clouds as partial input samples and the fusion of MLS and ALS point clouds as full tree references. This means that the partial point clouds exhibit real-world occlusion patterns. For both the partial and full trees, we first applied a statistical outlier filter, then computed the alpha shape and converted it back into a point cloud, as described in Section 2.2. We then infer the completion model inference on the MLS-based partial tree hulls.

To quantify the similarity between point clouds, we used the Chamfer distance (CD), which also serves as the loss function in our completion model. CD is defined as the sum of the average distances between each point in a predicted point cloud *P* and its nearest neighbour in a ground truth point cloud *G*:

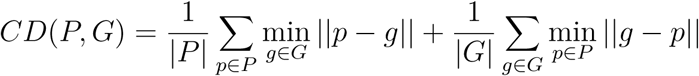

To compare regions where the predicted or partial crown differs most from the full reference, we also computed the Hausdorff distance (HD), which is defined as the maximum distance between any pair of nearest neighbours in the two point clouds (Taha and Hanbury, 2015):

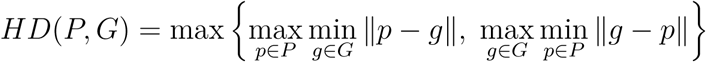

To evaluate the model’s crown shape predictions, we calculated both CD and HD between the full reference and the partial hull point clouds, as well as between the full reference and the predicted hulls. We used the Python library point-cloud-utils (Williams, 2022) to compute these metrics.

Additionally, we visualised the nearest neighbour distances on an example tree using cloud-to-cloud distance (C2CD) in CloudCompare software v2.13 (Girardeau-Montaut, 2019).

#### 2.4.2. Tree height prediction

To evaluate how accurately the completion model can predict actual tree height, we used the SWBM dataset. This dataset primarily consists of large trees (approx. mean height = 32 m) from dense conifer stands, and was acquired using a mid-range TLS device (Leica BLK360). As a result, it provides a representative example of situations where the upper part of the crown is missing from the point cloud, making it difficult to derive true tree height from ground-based LiDAR data alone.

Although full point clouds were not available for this dataset, we had access to tree height measurements from a field inventory, which served as the reference for actual tree height.

We downsampled each single-tree point cloud to a maximum of 100,000 points, then computed the alpha-shape and converted it back into a point cloud, following the procedure described in Section 2.2. We performed model inference ten times, using the ten models trained on different train/validation splits. Finally, we calculated the height of the predicted hulls as the extent in the z-dimension, and compared it to the field-measured tree height.

#### 2.4.3. Influence of proportion of missing tree height

To evaluate how much missing tree height the model can successfully reconstruct, we used the full trees from the pytreedb dataset (a fusion of TLS and ALS) because it consists of high-quality and complete TLS and ALS data. We generated full and partial hull point clouds using the same procedure as described in Section 2.2, downsampling each single-tree point cloud to a maximum of 100,000 points. However, we set a constant alpha value of 0.3 and, instead of removing a random percentage of tree height, we created test sets where we systematically removed 10%, 20%, 30% and 40% of the tree height.

## 3. Results and Discussion

### 3.1. Crown shape prediction

We evaluated the model’s ability to reconstruct full tree crown shapes using the hdALS-MLS dataset, which includes trees with missing lateral crown sections due to occlusion or segmentation errors. For this analysis, we compared both partial (MLS-based only) and predicted tree hulls against the full reference hulls using nearest neighbour-based distance metrics. Figures 3A–B illustrate the cloud-to-cloud distance (C2CD) for an example tree, visualising local differences between the full reference and both the partial and predicted hulls. The predicted crown exhibits generally smaller distances to the reference, indicating that the model successfully recovered much of the missing structure. Figures 3A-B summarise the distribution of Chamfer Distance (CD) and Hausdorff Distance (HD) for all 80 trees in the hdALS-MLS test set. On average, mean CD per tree decreased from 0.39 m (± 0.12 m) for the partial trees to 0.32 m (± 0.10 m) for the predicted trees. Similarly, mean HD decreased from 1.83 m (± 0.99 m) to 1.48 m (± 0.94 m). These results show that our completion model consistently improved the similarity between partial and full crown shapes, effectively reconstructing occluded or missing crown segments. Qualitatively, this is also visible in Figure 4, where the predicted alpha-shapes coincide well with the alpha-shape of the fused MLS and ALS tree point clouds. On the bottom right is an example where the model was unsuccessful in predicting the shape of the full tree. Notably, the model does not hallucinate any incorrect shapes but rather assumes that the tree is already complete.

**Figure 3:**
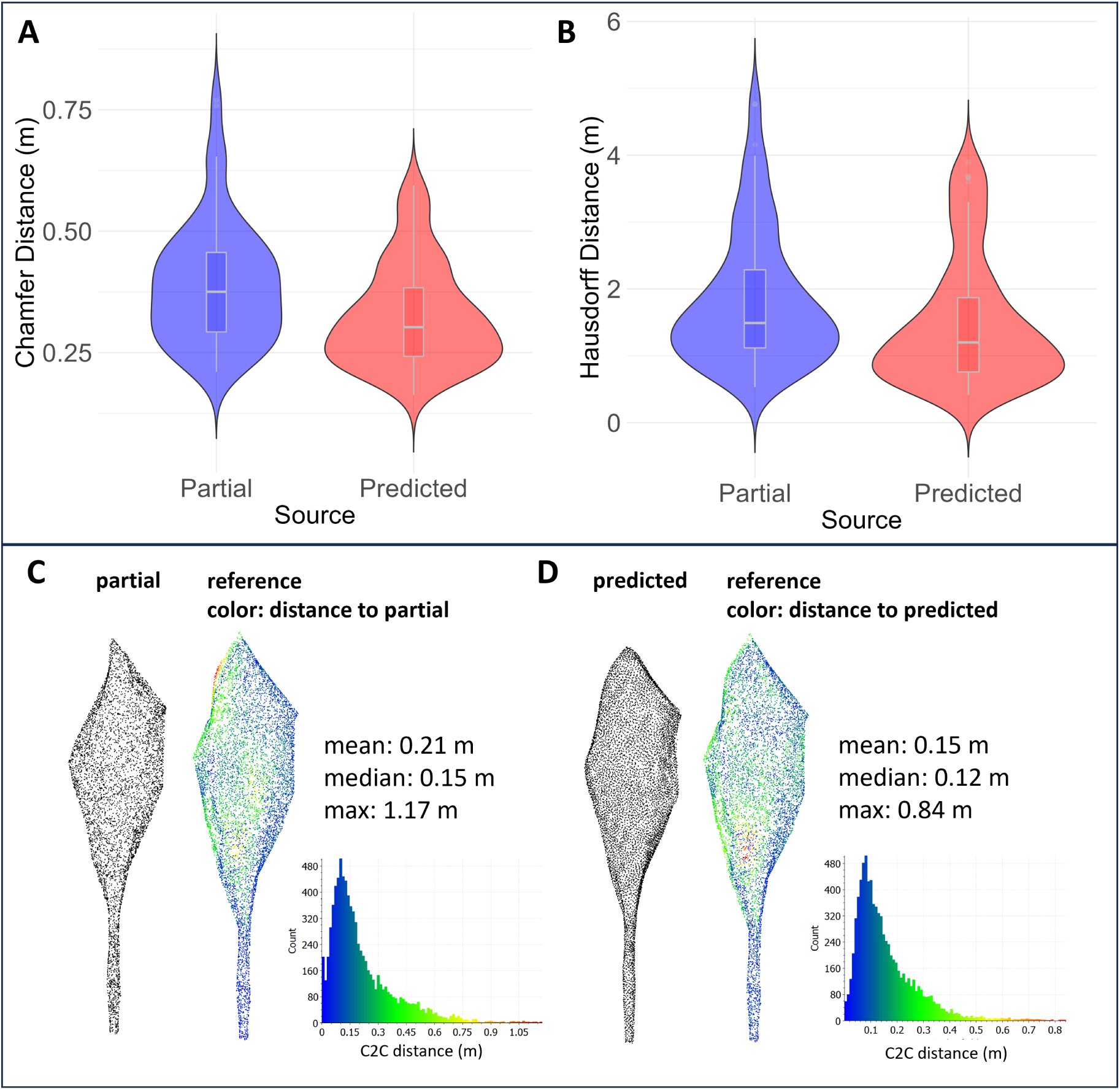
A-B: Chamfer Distance (CD) and Hausdorff Distance (HD) for partial vs. reference and predicted vs. reference tree hulls from the hdALS-MLS test dataset. C-D: Illustration of cloud-to-cloud distances (C2CD) of (C) partial vs. reference and (D) predicted vs. reference hull point clouds for one example tree.

**Figure 4:**
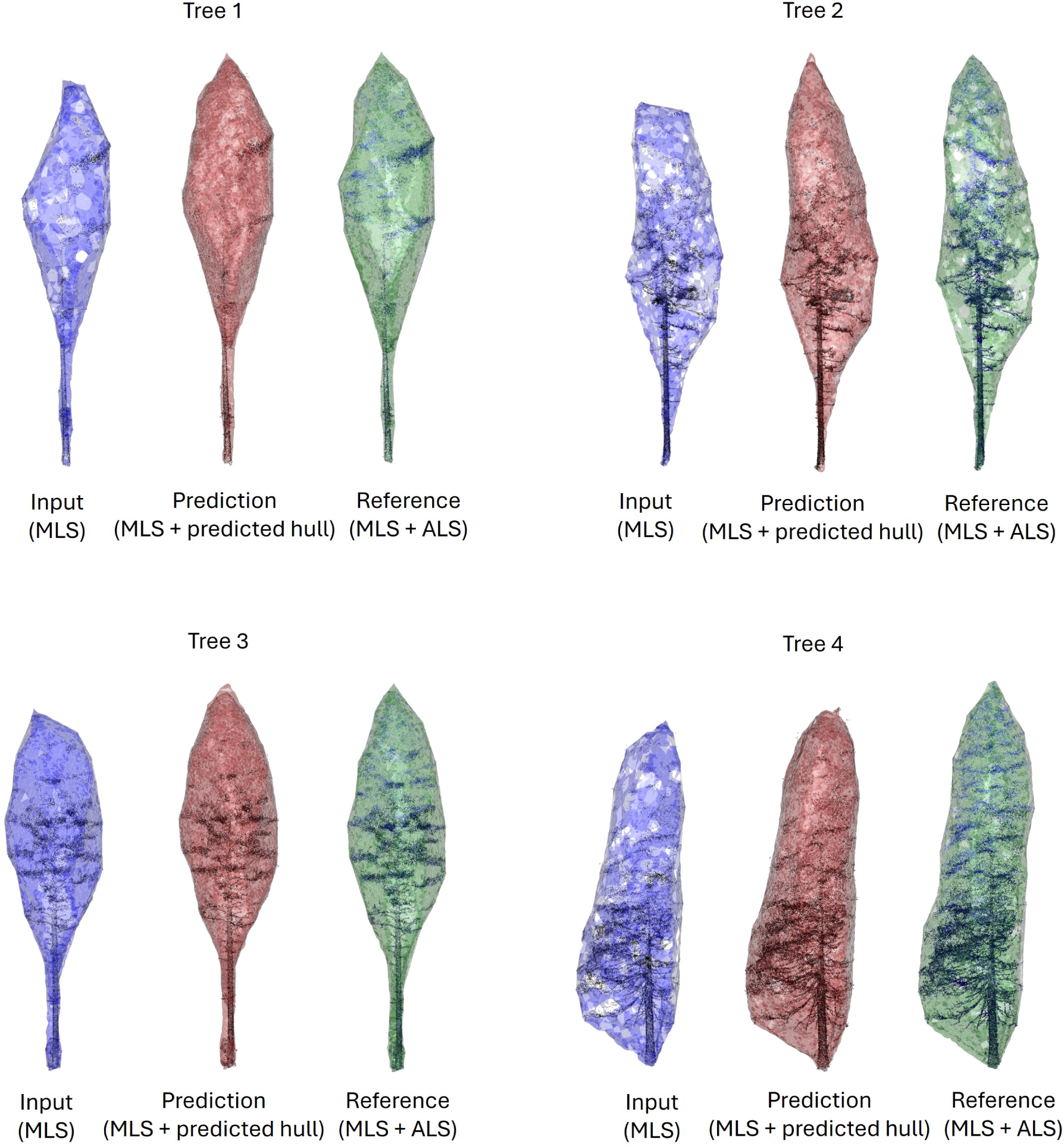
Four examples of completing tree crown alpha-shapes from the hdALS-MLS (NIBIO) test dataset: incomplete input tree crown shapes derived from MLS data are in blue, predicted alpha-shapes are in red, and reference alpha-shapes are in green.

### 3.2. Tree height prediction

Using the SWBM dataset, we evaluated the model’s ability to predict real tree height based on single tree point clouds with incomplete tree tops as acquired in the field using a Leica BLK360 terrestrial laser scanner. Figure 5A shows the distribution of field-measured tree height (reference) compared to the heights of both the input point clouds (RMSE = 5.04 m) and the predicted full tree hulls (RMSE = 3.68 m). The partial tree heights had a bias of -11.8% relative to the reference height, while the model predictions reduced this bias to -3.5%. Except for some outliers, the predicted tree hull heights agree well with the field measurements. However, it has to be stated that there is also considerable uncertainty in tree height measurement using hypsometer instruments (Stereńczak et al., 2019).

**Figure 5:**
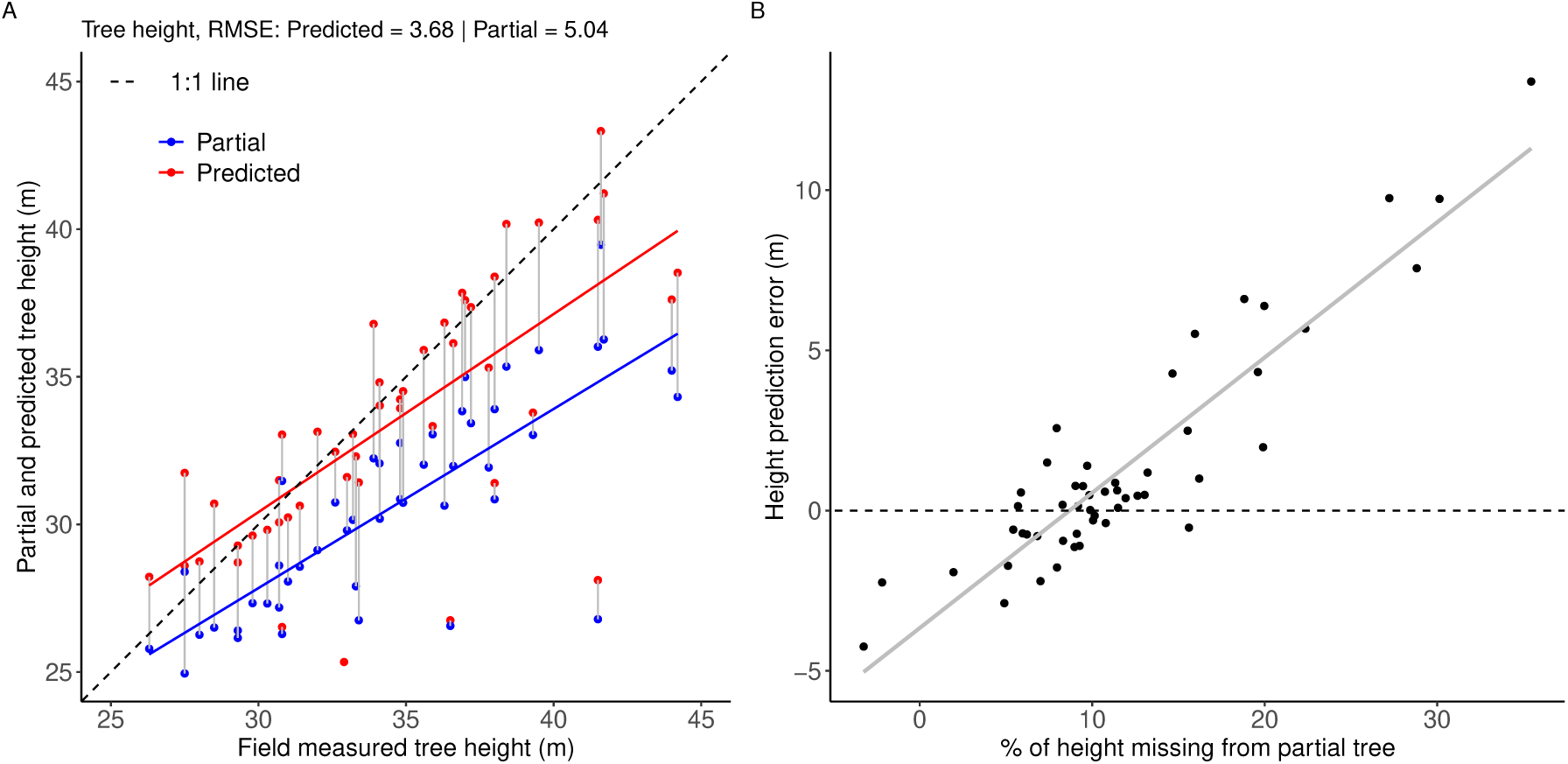
A) Height of the original partial (blue) and predicted tree hulls (red) vs. field-measured reference tree height. Partial and predicted values for the same tree are connected by gray lines. B) Percentage of tree height missing from the input point cloud vs. the difference between field-measured and predicted height.

A visual comparison of predicted tree hulls and field-measured tree heights is shown in Figure 6, illustrating four successful cases and one unsuccessful case. The last example illustrates a tree whose partial point cloud already resembles a full crown. This likely led the model to infer it was complete, and thus failed to predict the missing top segment. This tree corresponds to the outlier in the bottom right of Figure 5.

**Figure 6:**
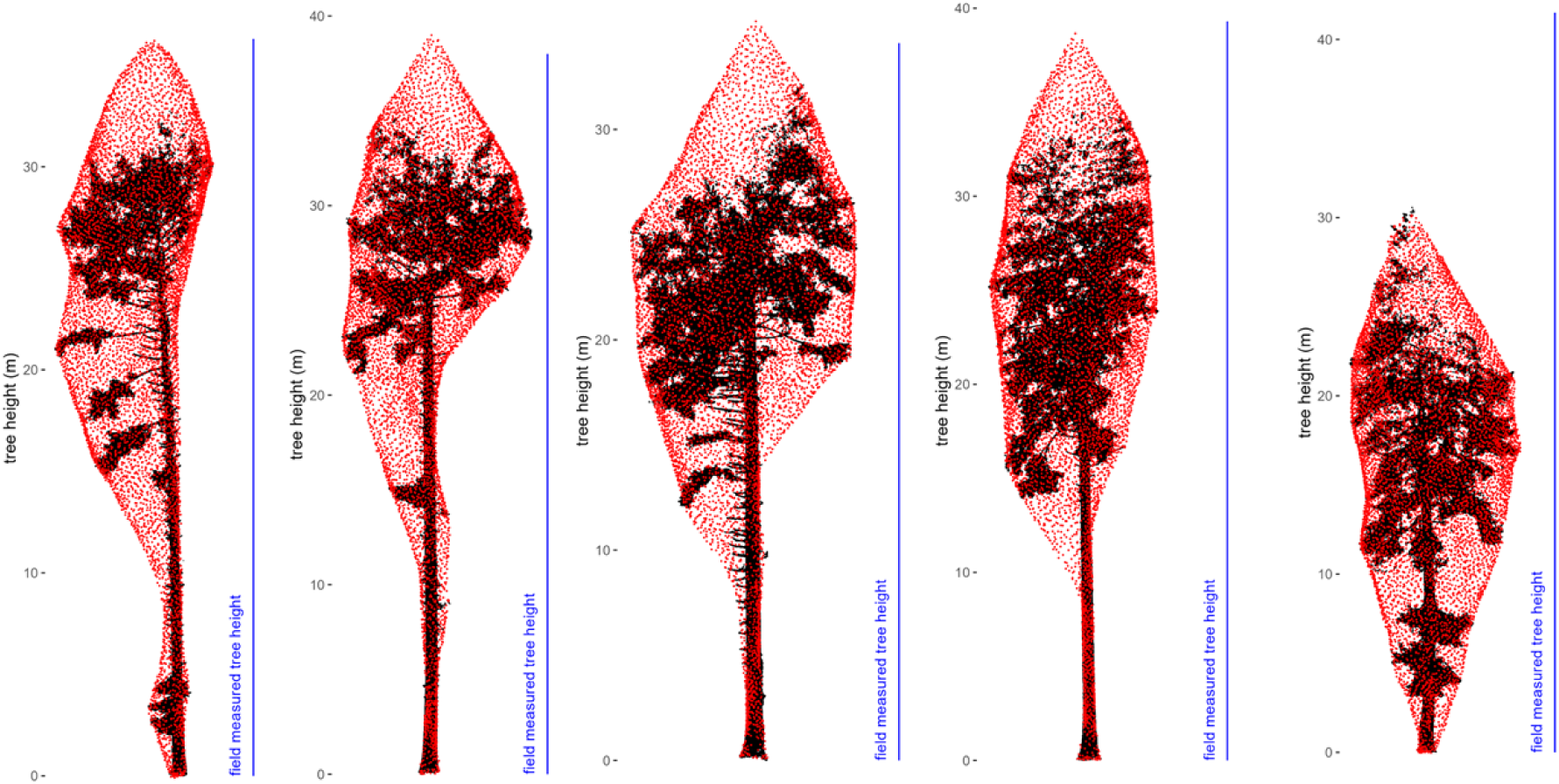
Examples of incomplete TLS point clouds (black), predicted hulls (red), and field-measured reference tree height (blue line).

Overall, we observe that the model tends to slightly overestimate height or, in some cases, fails to add any missing height at all. Given the way the training data were generated, this is reasonable. There is a high number of training samples that retained the full tree height, with only lateral crown sections removed (see Figure 1B). Because the point removal was done using simple spherical masks, the resulting partial point clouds often have smooth cut-offs. In contrast, real TLS and MLS scans typically show a gradual decrease in point density in the upper canopy. If even sparse points are present near the tree top, the resulting alpha-shape may still resemble a complete crown, which likely prevents the model from predicting additional height.

To better deal with such cases, future training datasets could simulate partial crowns using more realistic occlusion patterns. For instance, Ge et al. (2024) proposed simulating MLS data with non-uniform density loss inspired by electromagnetic wave attenuation, which may better mimic real-world occlusion and distance effects.

### 3.3. Influence of proportion of missing tree height

Using the pytreedb dataset, we assessed the model’s sensitivity to the amount of missing points at the tree top. Figure 7 shows the height of predicted tree hulls and the corresponding partial input where we removed 10%, 20%, 30% and 40 % of the tree height. For all of these four test sets, we observe a smaller RMSE for predicted than partial input. For 10% and 20% of height removed, the model is able to predict the original tree height very well. However, the model performs best for medium-sized trees, while for trees under 25 m, the model tends not to add any height in the prediction. This pattern likely reflects a bias in the training dataset toward medium-sized trees, with most samples falling between 25 and 35 m in height (Figure 1). Since both training and inference inputs are scaled and normalized, the effect is not directly caused by point cloud size. Rather, it may stem from the more uniform and representative crown shapes typical of medium-sized trees, which the model learns more effectively. In contrast, smaller and larger trees, which are less frequent in both the training and test sets, often display more variable structures, making them more difficult for the model to reconstruct accurately.

**Figure 7:**
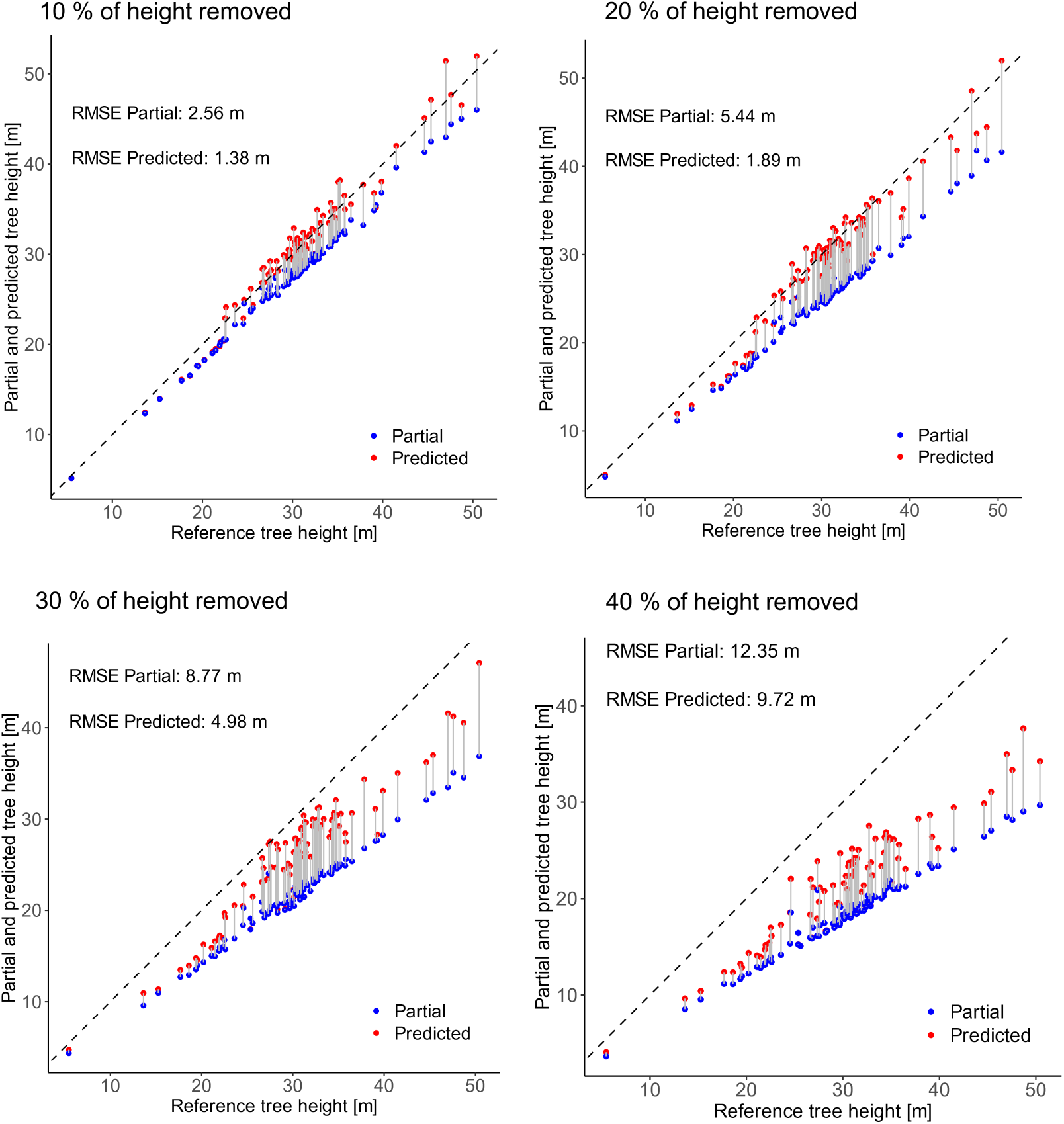
Height of partial (blue) and predicted tree hulls (red) vs. the full tree height. Results from four height thresholds with 10%, 20%, 30% and 40% of tree height removed from the tree top. Partial and predicted values for the same tree are connected by gray lines.

When 30% of the height is missing, the model systematically underestimates the true height. This effect becomes even more pronounced at 40% missing height. This is consistent with the model’s training regime, where a random proportion of 2% to 50% of the upper canopy was removed to simulate occlusion. As shown in the distribution of removed heights (Figure 1B), training samples with 20% height removal are more frequent than those with 30% or 40%, potentially explaining the model’s reduced performance in those higher-occlusion scenarios.

Figure 5B provides additional insight using TLS data from the SWBM dataset, where tree tops are naturally occluded. In this dataset, most trees lack approximately 10% of their height compared to field-measured values. For most of these cases, the completion model successfully predicted the full tree height. Similar to the artificially created test samples, the prediction error increases with the proportion of missing height in the point cloud, confirming that model performance declines as more of the upper crown is absent.

The amount of occlusion or unobserved tree structure in the upper canopy when using ground-based LiDAR depends heavily on the instrument, acquisition pattern, and especially on the forest plot’s structure, composition, and terrain. Moreover, relatively few studies have attempted to quantify occlusion to assess the 3D coverage of such datasets (Bienert et al., 2010; Kükenbrink et al., 2017; Brede et al., 2022a). Kükenbrink et al. (2025) performed occlusion mapping and analysis on handheld MLS data from national forest inventory plots. Across the 29 plots in their study, they observed an average occlusion fraction of 15% in the upper canopy (above 10 m), with one particularly challenging plot showing a total occlusion of 38% of canopy volume. Although such occlusion estimates cannot be directly translated into the height removal percentages used in this study, they offer a useful approximation of the typical degree of canopy loss in practice. These findings suggest that in real-world scenarios, it is rare to encounter trees with more than 30% of their height missing from point cloud data. Beyond this level, the data may become too incomplete to support reliable extrapolation of forest structure.

### 3.4. Advantages and limitations of tree hull completion based on alpha-shapes

The alpha-shape–based tree hull reconstruction offers several advantages over full single-tree point cloud completion approaches. By focusing on the outer shape rather than internal branch structures, this method should simplify the learning task, lowering computational requirements and potentially reducing the need for large, detailed training datasets. Key tree metrics (such as height, crown diameter, projected crown area, and crown volume) can be derived from the reconstructed hulls. If the lower stem is not occluded by branches, DBH and even full stem volume can also be estimated, provided tree height is predicted accurately. Additionally, because alpha-shapes are not affected by internal point cloud occlusion, they offer a more robust solution for estimating crown volume than voxel-based methods, which tend to underestimate the upper canopy due to sparse data inside the crown (Wang et al., 2024). This shape-based approach is also more sensor-agnostic, accommodating variations in data quality, acquisition platform, and tree morphology more effectively than methods that aim at reconstructing detailed internal structures.

However, completing only the outer hull instead of a full tree point cloud also comes with notable limitations. Most fundamentally, an alpha-shape provides no insight into the internal branch architecture of tree crowns and assumes a solid, continuous object, which oversimplifies the complex structure of the canopy. Computationally, alpha-shape generation is sensitive to point density and distribution. Sparse or uneven point clouds can lead to issues with triangulation or non-manifold shapes. Also, implementation differences across libraries can further affect robustness and consistency. Selecting an optimal alpha value is also not straightforward, since the appropriate value depends on the object’s scale, morphology, and the intended application (Gardiner et al., 2018). Alpha-shape volume is particularly sensitive to this parameter, especially for trees with many concave structures. Caution is therefore required when interpreting alpha-shape volumes derived from the predicted tree hulls, particularly if these volumes are intended for biomass estimation, as the associated uncertainties will require further investigation.

### 3.5. Outlook

This study introduced and evaluated a novel approach for reconstructing tree hulls from incomplete single-tree point clouds, with a focus on four coniferous species for which sufficient high-quality data were available. While the method shows promising results for conifers, future work should assess its applicability across a broader range of forest types and data sources. We expect that the alpha-shape-based approach can also be applied to broadleaf trees and could be relevant when scanning dense forest stands in leaf-on state. However, for broadleaf species, the branch structure and volume are often of interest. For such cases, models that predict the full point cloud would be more useful, especially those that integrate the skeleton structure into the model design (Xu et al., 2025; Zhang et al., 2025; Ge et al., 2024). If dealing with detailed point clouds with smaller gaps, a segment-wise completion approach (Bornand et al., 2024) could be used for broadleaf species with clear branching structures.

A possible downstream application of our proposed method could be to use the predicted tree hulls as spatial constraints for branch reconstruction models, such as the one proposed by Wang et al. (2023), where they used alpha-shapes as the outer boundary for a branch growth model. For coniferous trees, the method could also simply be combined with stem volume models by using the predicted tree height as an upper boundary for volume extrapolation.

While this study focused on completing the upper parts of the tree crown, it would also be interesting to train the model to complete any missing part of a tree, for example, the lower parts of tree stems in UAV-based data. This would be particularly relevant for cost-effective SfM point cloud data that usually does not “see” through the canopy as well as LiDAR sensors (McNicol et al., 2021). However, training and/or testing this approach on SfM data requires a dataset where full point clouds (preferably from TLS) are available in addition to the incomplete SfM data.

Generally, a key requirement for further development of tree completion models is the availability of suitable training data. However, evaluating the quality and completeness of training point clouds remains a challenge. Currently, this is done via visual inspection, but mapping of occlusion or unobserved space may offer a more systematic way to assess data completeness in the future (Gassilloud et al., 2025).

## 4. Conclusion

In this study, we explored the novel use of deep learning-based point cloud completion to reconstruct the outer crown shape (or hull) of individual coniferous trees from incomplete point cloud data. This approach is particularly relevant for forestry applications relying on ground-based remote sensing data such as TLS or MLS, where upper canopy occlusion is common and can hinder accurate tree metric extraction.

By targeting the crown hull, the model remains general and less susceptible to overfitting yet still delivers meaningful structural information. Across three diverse datasets, we demonstrate that the method improves both geometric similarity to full crowns and the accuracy of height estimation compared to using partial point clouds alone. The model performs robustly even under moderate occlusion and across varied acquisition conditions, reducing error in key metrics such as tree height, which are critical for forest inventory and ecological modelling.

This makes the method well-suited for practical use in forest monitoring workflows, particularly in conifer-dominated stands where the outer shape of the crown provides valuable indicators of tree growth, competition, or structural diversity. The ability to recover plausible crown geometry from incomplete scans can improve single-tree biomass estimation and serve as pre-processing for further structural analysis.

While performance declines when larger portions of the tree crown are missing, especially at the treetop, the model tends to avoid implausible reconstructions. Future work should focus on improving realism in training data to better reflect natural occlusion patterns and sensor-specific data artefacts, and explore the applicability to different species and forest types. Still, the approach offers an immediate, practical contribution to enhancing the usability of ground-based 3D data in remote sensing pipelines for forestry applications.

## Declaration of AI and AI-assisted technologies in the writing process

During the preparation of this work, the authors used OpenAI’s ChatGPT v4.0 to improve readability. After using this tool, the authors reviewed and edited the content as needed and take full responsibility for the content of the publication.

## Data availability

Code available at https://github.com/alBrnd/treePoinTr and the full public datasets of which parts were used in this study are available at https://doi.org/10.5281/zenodo.13255198 (FOR-species20K), https://doi.org/10.1594/PANGAEA.942856 (pytreedb), and https://www.doi.org/10.16904/envidat.403 (SWBM).

## Author contributions: CRediT

**A.B.**: Writing – original draft, Writing – review & editing, Formal analysis, Investigation, Data curation, Methodology, Conceptualization. **M.A.**: Writing – review & editing, Supervision, Funding acquisition. **F.M.**: Writing – review & editing, Supervision, Funding acquisition. **S.P.**: Data acquisition, Data curation, Writing – review & editing **R.A.**: Data acquisition, Data curation, Writing – review & editing **N.R**: Writing – review & editing, Methodology, Conceptualization, Supervision, Funding acquisition

## Appendix

### Model implementation details

For fine-tuning AdaPoinTr models on tree data, we mostly kept to the same implementation details and hyperparameters as described by Yu et al. (2023). They used AdamW optimizer to train the network with initial learning rate as 0.0001 and weight decay as 0.0005. The only change we made was to reduce the batch size to 16 and maximal number of epochs to 200. Training progress is shown in Figure 8.

**Figure 8:**
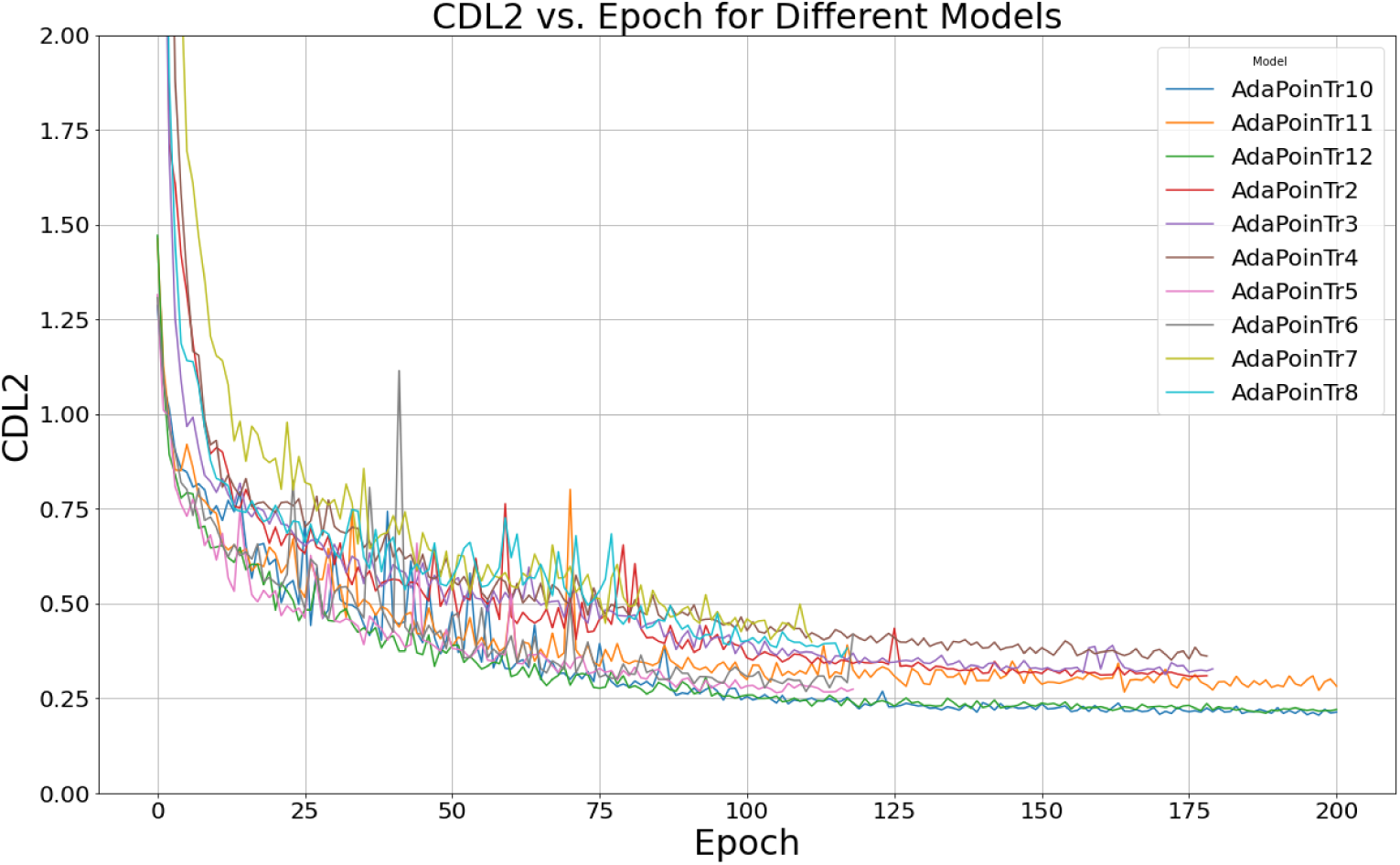
Training progress of 10 separate completion models with different (random) training/validation splits, but same hyperparameters. Chamfer distance (L2) was used as loss function.

### Additional Figure: Tree height prediction

We calculated the range of height predictions across ten model runs as a percentage of tree height, as shown in Figure 9. The average range was 4.9%, with a minimum of 0.7% and a maximum of 11.4%. This indicates some variation between models trained on different random train/validation splits, although the variability is relatively small.

**Figure 9:**
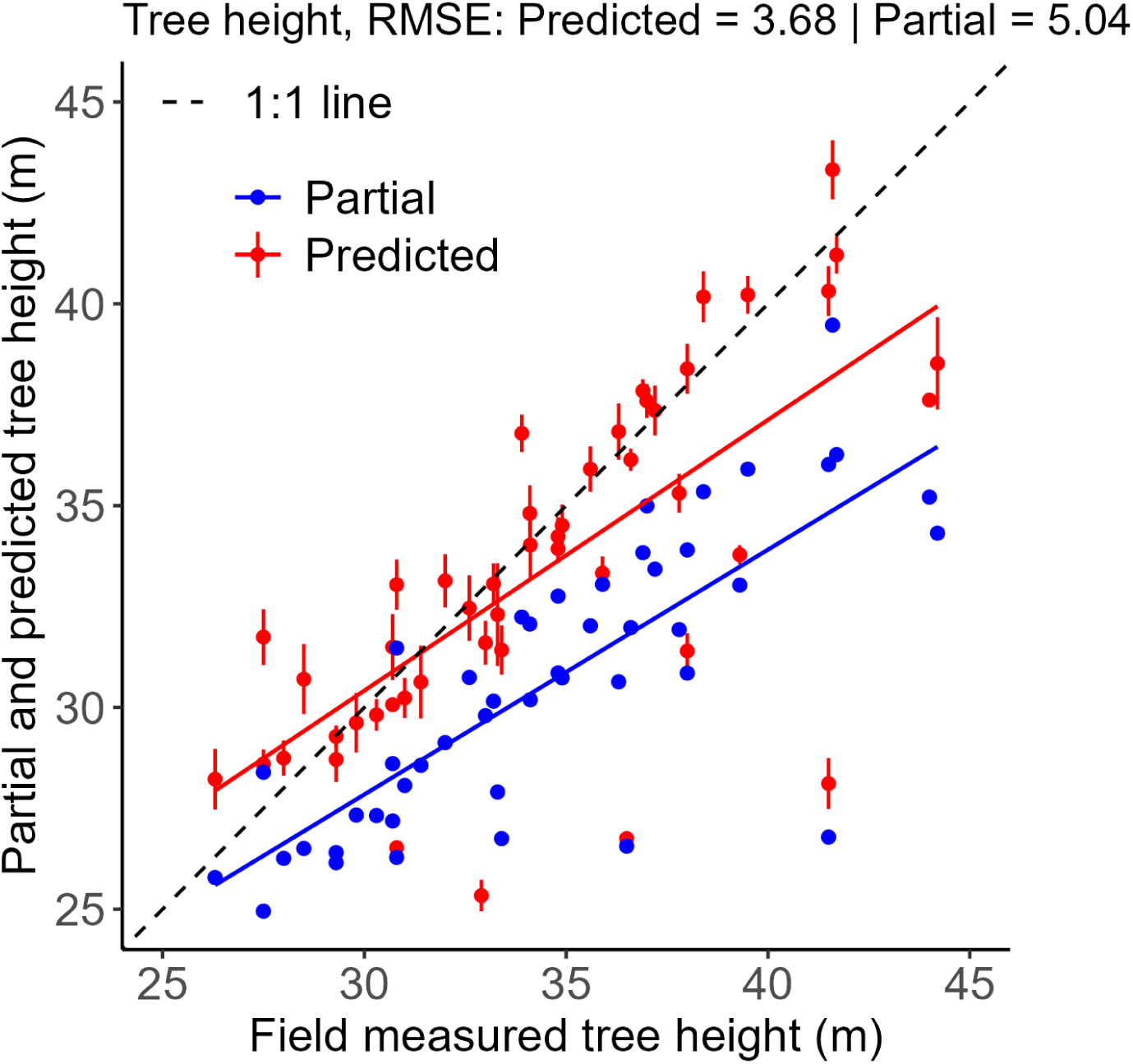
Height of partial (blue) and predicted tree hulls (red) vs. field-measured reference tree height. Error bars show the standard deviation from 10 completion models trained on different training/validation splits.

